# Genomic variant calling: Flexible tools and a diagnostic data set

**DOI:** 10.1101/027227

**Authors:** Michael Lawrence, Melanie A. Huntley, Eric Stawiski, Art Owen, Thomas D Wu, Leonard D Goldstein, Yi Cao, Jeremiah Degenhardt, Jason Young, Joseph Guillory, Sherry Heldens, Marlena Jackson, Somasekar Seshagiri, Robert Gentleman

## Abstract

The accurate identification of low-frequency variants in tumors remains an unsolved problem. To support characterization of the issues in a realistic setting, we have developed software tools and a reference dataset for diagnosing variant calling pipelines. The dataset contains millions of variants at frequencies ranging from 0.05 to 1.0. To generate the dataset, we performed whole-genome sequencing of a mixture of two Corriel cell lines, NA19240 and NA12878, the mothers of YRI (Y) and CEU (C) HapMap trios, respectively. The cells were mixed in three different proportions, 10Y/90C, 50Y/50C and 90Y/10C, in an effort to simulate the heterogeneity found in tumor samples. We sequenced three biological replicates for each mixture, yielding approximately 1.4 billion reads per mixture for an average of 64X coverage. Using the published genotypes as our reference, we evaluate the performance of a general variant calling algorithm, constructed as a demonstration of our flexible toolset, and make comparisons to a standard GATK pipeline. We estimate the overall FDR to be 0.028 and the FNR (when coverage exceeds 20X) to be 0.019 in the 50Y/50C mixture. Interestingly, even with these relatively well studied individuals, we predict over 475,000 new variants, validating in well-behaved coding regions at a rate of 0.97, that were not included in the published genotypes.

## 2 Introduction

A genomic variant is an observed nucleotide difference from some reference at a specific position in the genome. Detecting genomic variants at low allele frequencies is important for cancer research, and knowledge derived from variant calls drives the development of new therapies and diagnostics. Variant calling is distinct from genotyping, which requires an assumption of ploidy and an assumption that the cells being sequenced share a common genome. In the diploid case, we expect nucleotide frequencies at 0, 0.5 or 1 [DePristo et al., 2011], [Li et al., 2009]. Genotyping of diploid organisms is straightforward and the concordance between different methods is high provided there is sufficient coverage. However, in tumors, variants have a wide range of allelic frequencies, in large part due to copy number changes and sample heterogeneity [Zhao et al., 2014]. Detecting variants in tumors remains problematic and there is limited agreement between methods [O’Rawe et al., 2013]. We believe that a primary reason for the discordance is the lack of an extensive data set with millions of variants, in different genomic contexts with approximately known frequencies. Such a data set would allow for direct comparison of methods across the entire genome.

There are many variant calling algorithms available, and indeed a variety of distinct objectives that are collectively described as variant calling (see Methods). For example, Varscan2 generates variant calls from the output of `samtools mpileup` [Koboldt et al., 2012], [Li et al., 2009]. It implements a number of tunable filters that consider variant frequency; read depths; biases in strand, read position and quality; and other aspects of the data. In contrast, LoFreq calls variants based on a Poisson-Binomial model that incorporates base quality information [Wilm et al., 2012]. Others have implemented error correction models [Wong et al., 2014]. However, there is little concordance among current callers and the concordance decreases when low frequency variants are considered [O’Rawe et al., 2013]. A flexible and efficient set of tools suitable for generating and analyzing variant calls will facilitate the diagnosis and comparison of existing methods and help drive innovations in this field.

We present two contributions that aim to foster the development of better variant calling algorithms. The first is an experimentally-generated whole genome sequencing dataset, with a large number of known variants at a range of allele frequencies and located in a variety of genomic contexts. Algorithm developers can use the dataset as a reference when diagnosing algorithm performance. We have also developed a Bioconductor package, VariantTools, which is a flexible framework for manipulating and filtering variant calls during algorithm development and *adhoc* analyses. We apply VariantTools to the reference dataset to demonstrate how one might use it to generate a general, exploratory set of variant calls.

## 3 Methods

### 3.1 WGS of a CEU/YRI mixture

A good reference data set should be based on real data and generated by applying current tools and protocols so that it reflects error rates that users are likely to see in practice. Further, it should have large numbers of well-characterized true positive variants that are present in different genomic contexts and at varying frequencies. Genomic context is important, because it affects the accuracy of both sequencing and alignment, as well as other aspects of the experiment. Li [Li, 2014] recognized the need for real data in diagnosing variant calling algorithms and sequenced a haploid cell line, where heterozygous calls were reasonably assumed to be errors. We complement his work by performing whole genome sequencing on a series of titrated cell mixtures to evaluate caller performance on heterogeneous samples.

Others have sequenced titrated cell mixtures over targeted, well-behaved regions [Stead et al., 2013], but aligners struggle to find unambiguous alignments in low complexity regions, or in regions with strong homology to other parts of the genome. The sequencing error rate depends on factors like the local GC content and the presence of homopolymers.

We mixed DNA extracted from two Coriell cell lines, NA12878 and NA19240, corresponding to the mothers of the HapMap CEU and YRI trios (GQ12878 and GQ19240, respectively). To achieve a broad range of frequencies, we mixed these reagents at ratios of 10Y/90C, 50Y/50C and 90Y/10C, in triplicate, (Figure 1A). This is expected to generate frequencies at 5%, 10%, 45%, 50%, 55%, 90%, 95% and 100% in the 90/10 mixtures, and 25%, 50%, 75% and 100% in the 50/50 mixture (Figure 1B). While variants can be present at <5% in complex samples, 5% is a reasonable lower bound, given the typical coverage of a WGS experiment. Figure 1D shows how much variability there was between the replicates.

**Figure 1:**
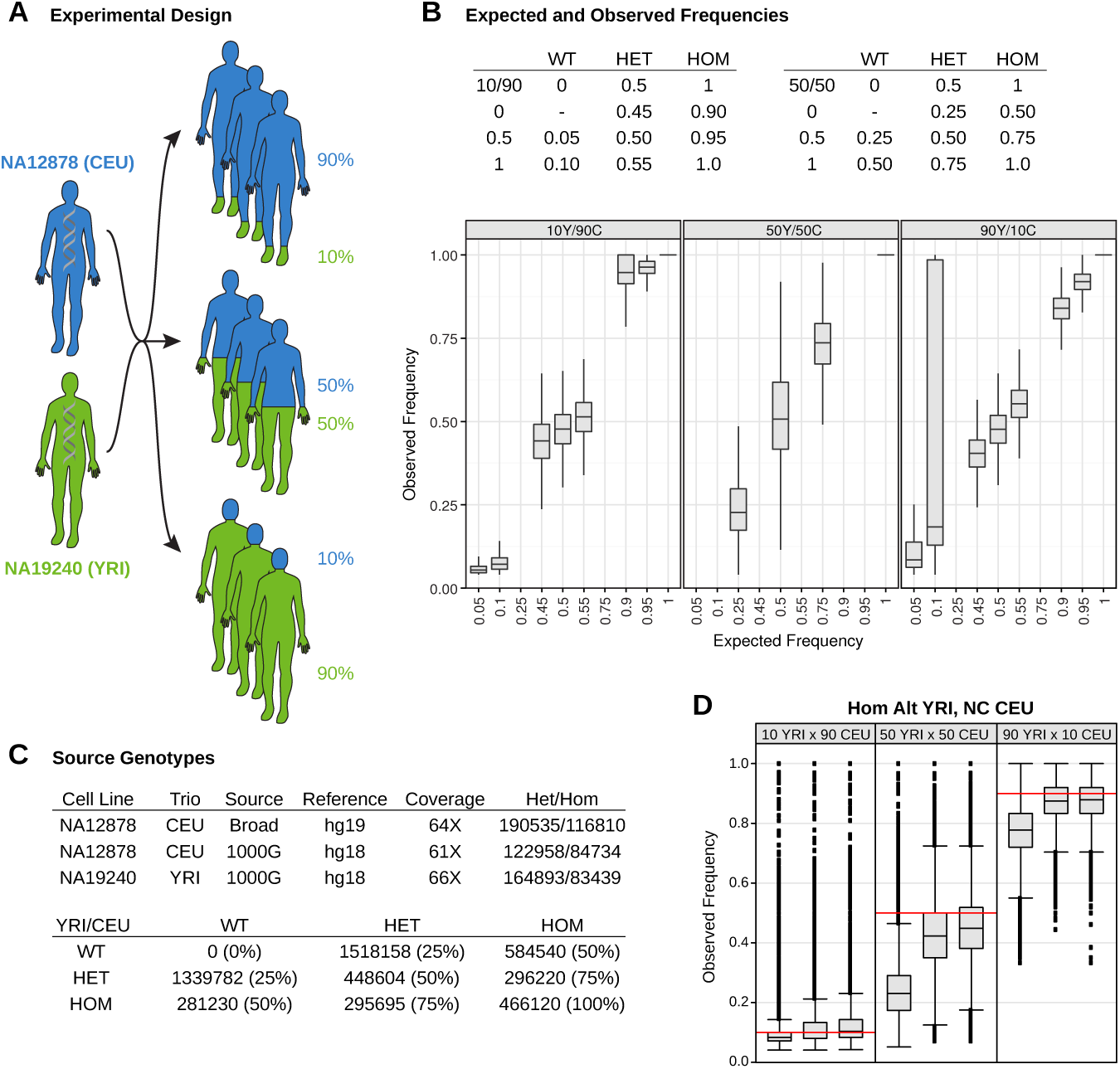
*Experiment.* (A) DNA from the NA12878 (CEU) and NA19240 (YRI) cell lines were mixed in three different proportions (10Y/90C, 50Y/50C and 90Y/10C), in biological triplicate. (B) Comparison of the expected and observed frequencies. The tables show the expected frequencies as a combination of mixture ratio and the genotypes of the two individuals. The boxplot shows the distribution of observed frequencies, conditional on the expected frequency and mixture. We suspect that the unexpectedly high frequencies for the 5% and 10% variants in 90Y10C are due to undercalling in the YRI. (C) Top, the source of the individual reference genotypes. Bottom, the counts and frequencies for variants in the 50Y50C (D) The observed frequency distribution in each triplicate for variants called homozygous alt in YRI but WT in CEU. The red line corresponds to the expected frequency if our mixture of the two DNA sources was perfect. Several of the samples, in particular the first replicate in each of the mixture ratios, are biased against YRI. This will yield lower observed variant frequencies and a corresponding lack of sensitivity in the YRI, especially in the 10Y90C mixture.

### 3.2 Reference genotypes

The two individuals have been sequenced and genotyped by a variety of approaches [The 1000 Genomes Project Consortium, 2012], [DePristo et al., 2011], and NA12878 is the subject of the Genome in a Bottle Consortium [Genome in a Bottle Consortium, 2014]. We obtained SNV calls for both from the 1000 Genomes Pilot 2 study, which incorporated sequencing from three different technologies (Illumina GA II, ABI SOLiD, and Roche 454) at a total coverage of 66X. These calls were lifted over from hg18 to hg19 using the `liftOver` function from the *rtracklayer* package [Lawrence et al., 2009]. We extracted an additional set of NA12878 SNV calls from the GATK Resource Bundle. The calls were generated by GATK, according to best practices, from a 64X Illumina HiSeq whole-genome sequencing dataset, aligned with BWA to hg19 (Figure 1C).

The 1000G calls are based on older tools and technology compared to those from GATK, and the 1000 Genomes genotype calls were made in a way that controls the false positive rate and hence there is a relatively high false negative rate for the 90Y/10C mixture (Figure 1B). This impacts our estimates of the FDR, as many of the supposed false discoveries may in fact correspond to true positives that are present in the genomes, but were not included in the reference genotype.

We first merged the 1000G and Broad CEU genotypes, taking the union of the two call sets to form a single CEU genotype. Where both sources called the same alternative allele, we took the het/hom determination from the Broad genotypes. We then constructed a reference genotype for the mixtures by combining the YRI and CEU genotypes into a single reference in the same way. We allowed for multiple different alternative alleles at any locus. The variants present at each locus are constant across mixtures, but the frequency of the variant in each mixture depends on the genome of origin and the proportion of that genome in the mixture. The combined reference genome for the 50-50 mixture is described in Figure 1C.

We identified a specific set of variants, which we refer to as the *YRI-specific variants.* These variants were homozygous non-reference in the YRI genotype and were not called as variant in the CEU genome (and hence were presumptive homozygous reference). Thus their frequencies should track at 10%, 50% and 90% across our mixtures 10Y/90C, 50Y/50C and 90Y/10C.

### 3.3 Variant calling with GSNAP and VariantTools

We have developed the VariantTools Bioconductor package as a modular toolkit for generating, filtering and comparing sets of variant calls [Lawrence et al., 2014]. Users can construct variant calling pipelines by combining filters, including those defined by the user. Integration with Bioconductor facilitates annotation of variants and analysis of the association between variants and their genomic context. We demonstrate an application to the latter in Section S11.

For the sake of comparing a general variant calling approach to one based on the standard BWA/GATK workflow, we applied *VariantTools* to construct a simple, example variant calling pipeline, as diagrammed in Figure 2A. We aligned the reads using GSNAP [Wu and Nacu, 2010] (see S1.1 for the details) and tallied the nucleotides at each position, excluding base calls with a quality score of 23 or less. To focus this analysis, we restrict to SNVs, but indels are present in the data, and should be useful for the development of indel callers.

**Figure 2:**
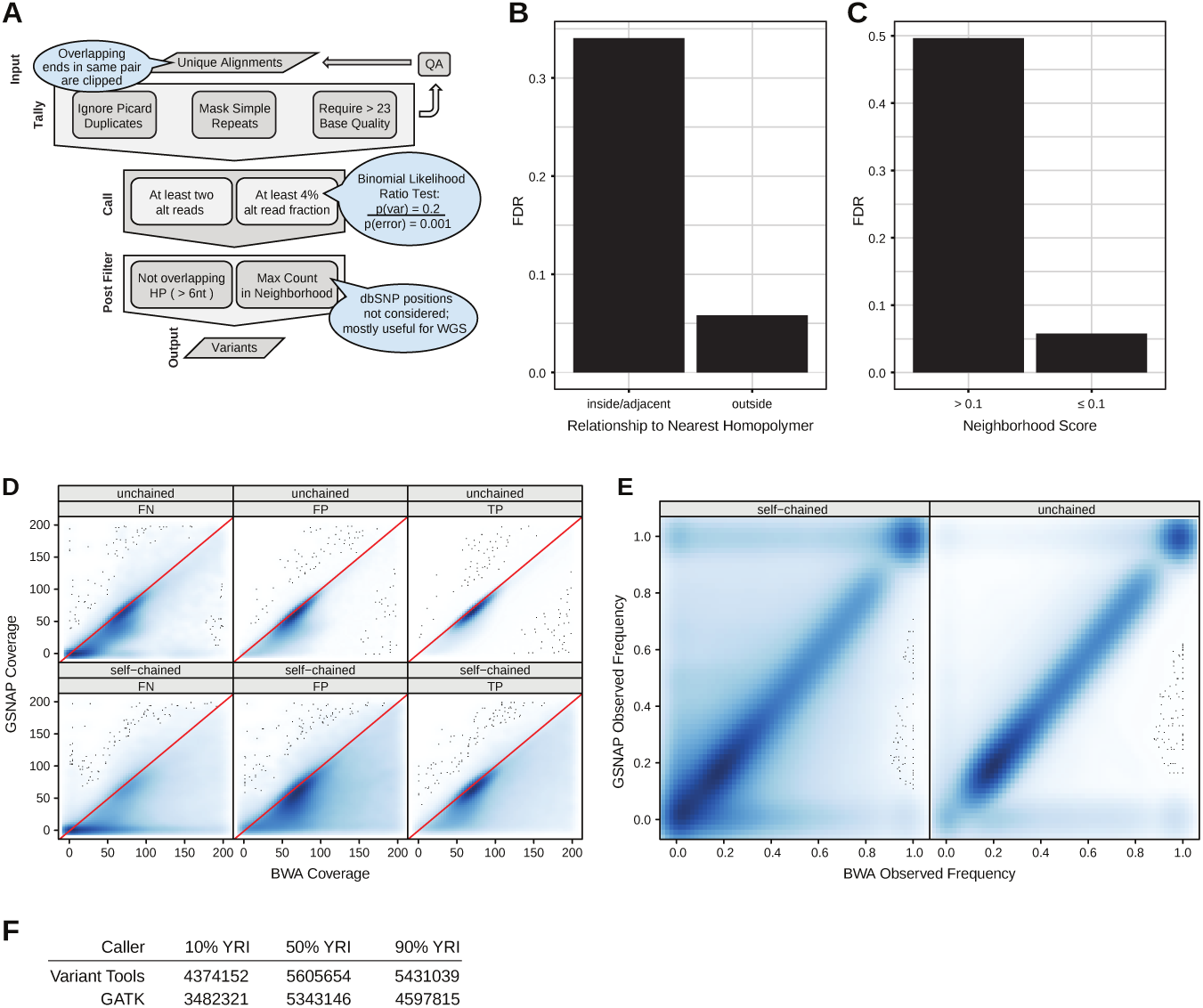
*Variant calling pipelines.* (A) Our workflow, based on GSNAP/VariantTools, converting the unique read alignments to variant calls by tallying the nucleotides and applying filters to the tallies. (B) The FDR (see Methods) is higher for variants over homopolymers of length > 6. (C) The FDR is higher for variants with a neighborhood score (see Methods) above our filter cutoff of 0.1. (D) Comparison of the BWA and GSNAP coverage at the union of the published alt genotypes and VariantTools calls, conditional on the concordance in terms of false negatives (FN), false positives (FP) and true positives (TP) and self-chain status (see Methods). BWA tends to have greater coverage, especially in self-chained regions, and there are many FN regions with moderate BWA coverage but near zero coverage in GSNAP. (E) Comparison of the BWA and GSNAP observed alt frequencies at alt positions in the published genotypes. Agreement is generally good. (F) The number of variants called by Variant-Tools and GATK, conditioned on the mixture. VariantTools called significantly more variants than GATK, especially in the imbalanced mixtures (10Y90C, 90Y10C), where variants were present at low frequencies.

Within the VariantTools framework, we defined a filter based on a likelihood ratio (equivalently a Bayes factor) as the basis of our method, along with several other filters (see S1.2 for details). We use a Binomial distribution to model the observed counts and use *p_E_* = 0.001 for the error rate with *p_V_* = 0.20 as a lower bound for the frequency of a true variant. The error rate and lower frequency bound are parameters and users will set them according to their needs. We chose 0.2 as the lower bound, which is slightly below the expected frequency of heterozygote variants in the 50-50 mixture and should give good sensitivity and specificity for that data set, which is our focus. We declare either error or variant, according to which probability is the larger. With our selected values of *p_E_* and *p_V_*, the test is equivalent to calling a variant when the observed non-reference frequency cutoff is above 0.04.

Others have reported extra Binomial variation [Plagnol et al., 2012], [Gerstung et al., 2012] and hence suggested alternatives, such as the Beta-binomial. We found (see Section S4) that extra-binomial variation is associated with whether the variant has a positive self-chain score, suggesting that it arises due to mapping/alignment issues.

### 3.4 Variant calling with BWA/GATK

For comparison purposes, we called variants with a pipeline based on commonly used tools. Reads were mapped to the UCSC human genome (GRCh37/hg19) using BWA [Li and Durbin, 2009]. We called variants with the GATK UnifiedGenotyper [DePristo et al., 2011] (see S2 for more details on the GATK pipeline). While other approaches exist, this pipeline meets our need for a baseline genotyping pipeline.

### 3.5 UCSC self-chain scores

We rely on the UCSC self-chain scores [Karolchik et al., 2014] for measuring similarity between different genomic regions (see Section S3). The self-chain score is a generic and robust indicator of intragenomic similarity that is independent of the aligners we used. This is in contrast to the mapping quality (MAPQ in the SAM spec), which represents the probability that an alignment is correct, under assumptions specific to the aligner.

### 3.6 Cell culture and sequencing

Three independent pairs of plates of NA12878 and NA19240 cells were cultured to 80% confluence in RPMI supplemented with 10% Fetal Bovine Serum, 1% Sodium Pyruvate and 1% Glutamine. Total DNA from these 6 plates was isolated with DNeasy Blood & Tissue Kit from Qiagen, and quantified by picogreen. Samples were mixed as described previously (Figure 1A). The 9 mixed samples were prepared for whole genome sequencing using the Illumina Truseq DNA sample preparation kit, and sequenced on an Illumina HiSeq.

## 4 Results

We generated a diagnostic dataset for variant calling by whole genome sequencing a titrated mixture series of two HapMap individuals (NA12878/CEU, NA19240/YRI) with well characterized genotypes. There were three mixture ratios (10Y90C, 50Y50C, 90Y10C), with three replicates each, and we merged the replicates to yield about 64X coverage per sample. We called variants using a custom general variant calling pipeline, as well as a best practice GATK pipeline.

### 4.1 Comparison of Alignment Pipelines

The two pipelines were in high agreement. They called variants at the same loci and the frequencies of the observed variants were highly concordant (Figure S7). However, regions of homology are challenging for aligners and in these regions the two approaches are more discrepant. In our hands BWA tends to uniquely align more reads at some loci than GSNAP, as shown in Figure 2D. The plot is divided by the GSNAP/VariantTools concordance versus the published genotypes in terms of false negatives (FN), false positives (FP) and true positives (TP). In general, the agreement is better in regions that are not self-chained. Figure 2E compares the BWA and GSNAP observed variant frequencies for the YRI-specific variants. We see strong agreement, although the agreement is weaker in self-chained regions.

Figure 2F shows that our general variant caller always detects more variants than GATK genotyper and that the differences are most pronounced when the mixture is imbalanced with one sample present at 10%.

### 4.2 Observed variant frequencies

Figure 1B suggests that the reference genotype is not as accurate as we had anticipated. While the boxplots for the 10Y/90C mixture look as expected, the observed frequencies of the 10% variants in the 90Y/10C do not. There are many high frequencies observed which would be consistent with real YRI variants missing from the reference genotype. We anticipated that the CEU genotype would be better estimated, since we derived it from two external sources, including newer data from the GATK project. The high frequencies are also consistent with the notion that the 1000G calling was conservative and strongly controlled the FP rate at the cost of an increased FN rate. Indeed, if we consider the variants with an observed frequency above 0.2 and an expected frequency of 0.1 in the 90Y10C sample, we find that 89% of them follow a frequency trend consistent with their being homozygous in CEU and non-ref in YRI (see S9 for details on the algorithm).

Accurate frequency estimation is important for characterizing the composition and evolution of tumors. Figure 3A presents the variant frequency distributions for the YRI-specific variants. The frequency peaks observed in Figure 3A match well with the mixture frequencies we achieved. While the frequency distributions for the nominal 5 and 10% variants overlap substantially, all others are well separated.

**Figure 3:**
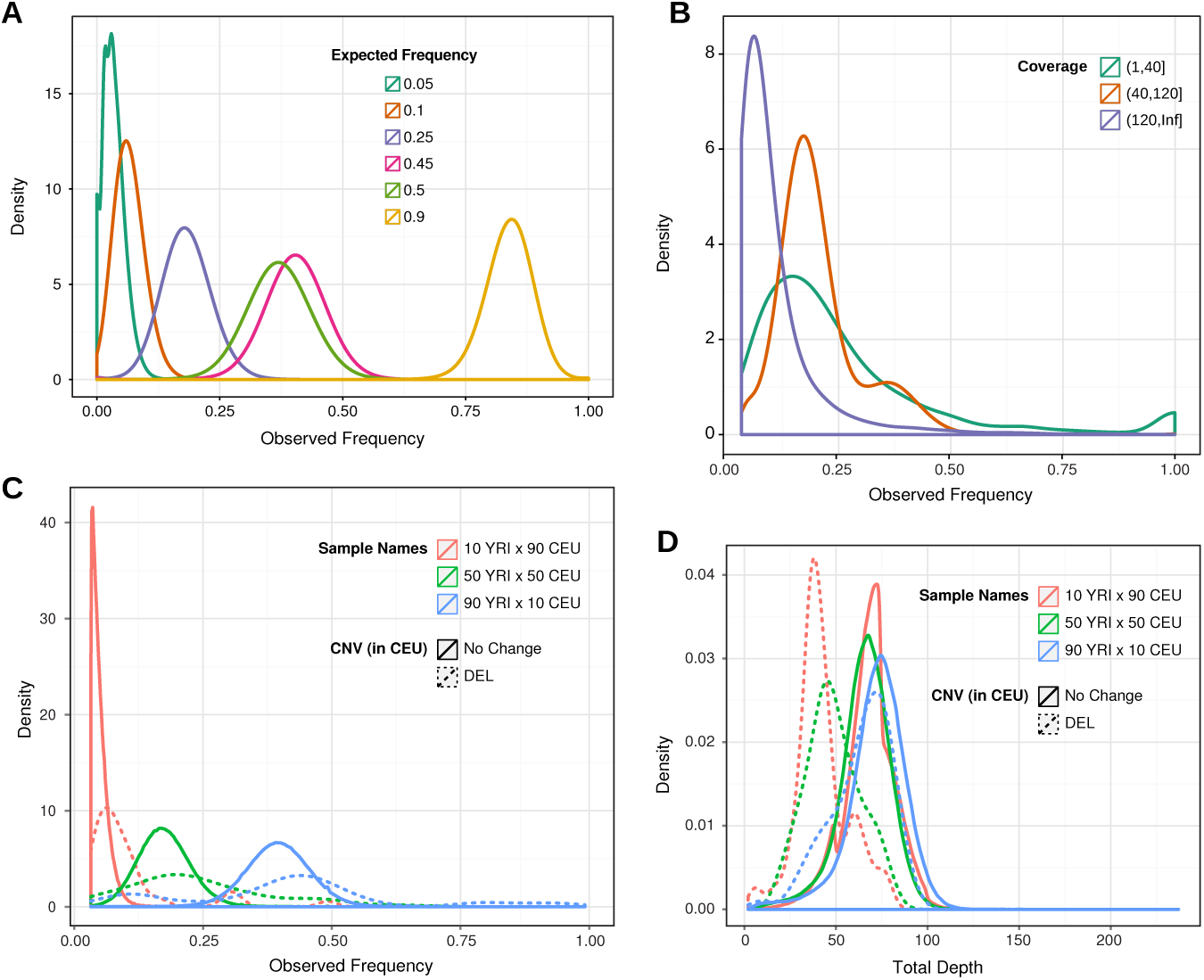
*Variant frequencies.* (A) Observed frequency distribution, conditioned on expected frequency, for the YRI-specific variants. We see reasonable separation and can distinguish 25% variants from those at 50%. The variants expected at 50% have a lower average frequency than those expected at 45%, which is due to the bias in the mixtures, see Figure 1 and Section 3.2. (B) Observed frequency distribution for the YRI-specific variants in the 50Y/50C mixture, conditioned on low (<= 40X), moderate and high (>120X) coverage. Due to unintended mixture bias against YRI, we should observe a bimodal distribution of frequencies at 20% and 40%. For moderate coverage that is the case, but in low or high coverage regions the frequencies do not show this pattern. (C and D) Distribution of frequency and coverage, respectively, for the YRI-specific het variants, conditioned on whether a variant falls within a CEU deletion. In C, the frequency distribution for CEU deletions has a higher variance and slightly higher mean than the overall distribution. In D, coverage within deletions appears to be lower on average.

Accurately estimating frequency depends on the coverage. Figure 3B shows that the distribution of YRI-specific variant frequencies in the 50Y/50C mixture, conditioned on coverage range. Only the frequency distribution for the moderate coverage variants aligns with our expected frequencies.

### 4.3 Concordance of variant calls

We now consider how concordant our calls are with the reference genotypes we constructed in Section 3.2. In Figure 4A we show the observed frequencies for variants across mixtures where variants are classified according to the known genotypes. We encode the genotypes according to YRI/CEU, where YRI and CEU can be either 0 (no-call), 0.5 (het) or 1.0 (hom alt). For example, a 1/0 variant is homozygous alt in the YRI but uncalled in the CEU. Such a variant should have an observed frequency of approximately 10% in the 10Y/90C mixture, increasing to 50% in the 50Y/50C mixture and 90% in the 90Y/10C mixture. The frequency trends largely match our expectations, although in the top-middle panel of 4A (CEU het, YRI WT), there is some evidence of error. Some loci have been misgenotyped due to undercalling in YRI, while others appear to be false positives in the CEU calls.

**Figure 4:**
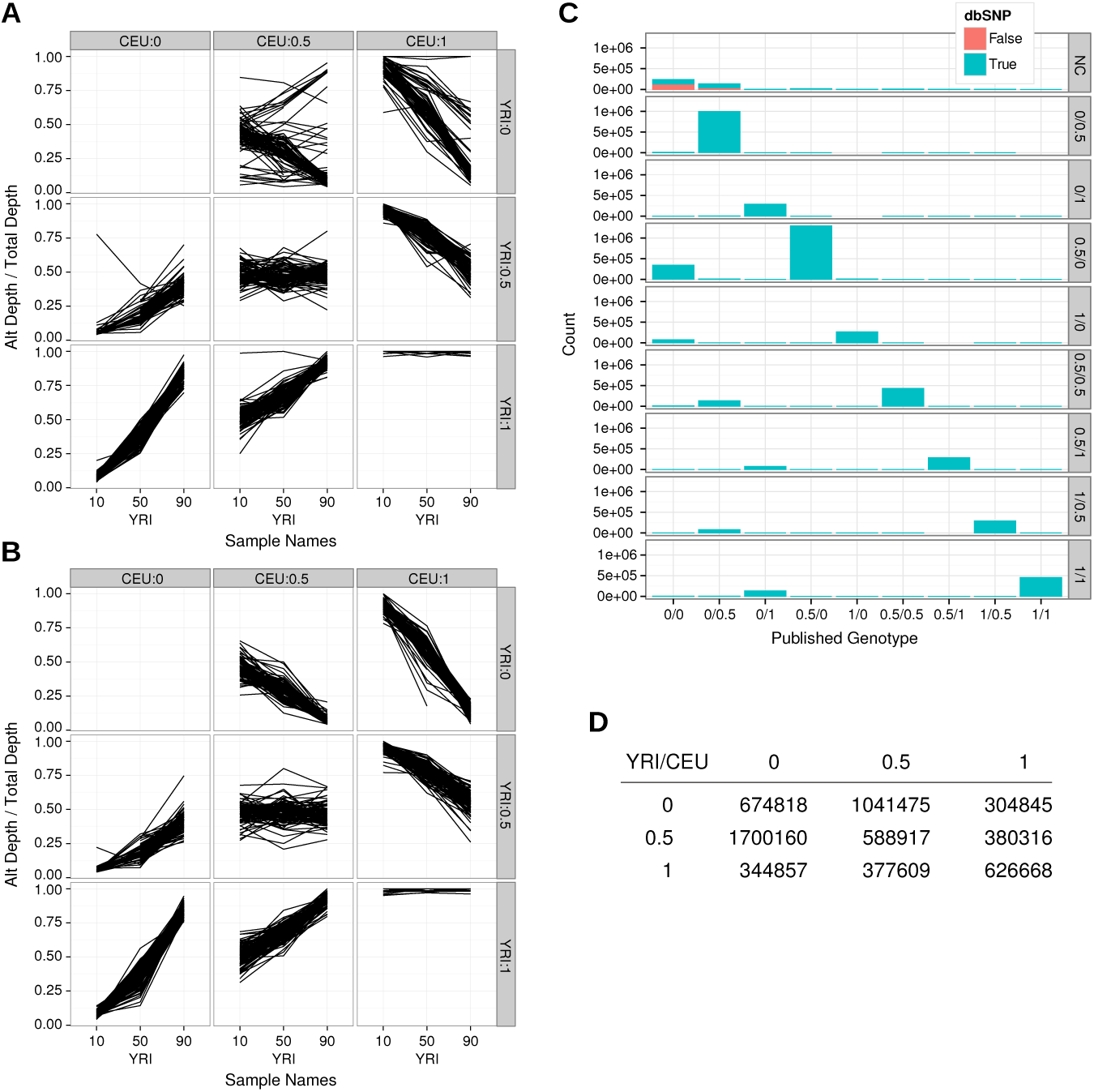
*Concordance of frequency trends.* (A) The frequency trends for a random subsample of 100 TP variants for each published genotype. Each line corresponds to a single locus. For each mixture, the observed frequency of the variant is computed and those points are joined by a line. (B) The frequency trends for a random subsample of 100 putative FP variants for each predicted genotype (see supplement). The trends in panels (A) and (B) appear quite similar, suggesting that putative FPs with predicted genotypes are likely to be TPs. (C) Cross-tabulation of the predicted and published genotypes, counting every unique variant called in any of the three mixtures. The color indicates presence in dbSNP. There is strong agreement, and most discordance reinforces our claim that the YRI genotype is undercalled. Those positions that are no-call (NC) in both the predicted and published genotypes are likely to be FP, which is consistent with their absence from dbSNP. (D) Count of each predicted genotype combination.

Due to the design of our experiment, the observed frequencies at any TP locus should track according to one of the patterns in Figure 4A. We exploit that fact to construct a genotype classification method for all variants (see S9). In short, we binned the observed frequencies and mapped particular sequences of bins to a genotype. For example, if a variant was observed at 10%, 50% and 90% for corresponding mixtures of 10Y/90C, 50Y/50C and 90Y/10C, we assign the genotype 1/0 (YRI: hom-alt, CEU: WT). We refer to a variant as “tracking” if its cross-mixture frequency pattern tracked an expected pattern and we predicted a genotype based on that pattern. Non-tracking variants were not assigned a predicted genotype and were labeled as No Call (NC). Figure 4D shows the count of each predicted genotype combination.

In Figure 4B we show the observed frequencies for a subsample of variants from each predicted genotype. While there is a strong similarity between Figures 4A and 4B, there are some discrepancies in the 0/0.5 panel of Figure 4A, which we suspected are due to missed YRI calls in the reference genotypes. We confirmed this with Sequenom-based validation (see Section 4.7).

We see strong concordance between the reference genotypes and our predictions in Figure 4C. The largest discrepancies are consistent with the observation that the YRI genome was substantially under-called in the reference. We found that 70% of the FP variants that we predicted to be in the YRI were excluded by the mask applied by the 1000G during calling (see Section S8). About 97% of those were masked due to having over 20% of the reads with a mapping quality of zero, which suggests that multi-mapping of the shorter, 36bp 1000G reads may have been a major factor in the lack of sensitivity.

Variant calls with a NC status in the predicted genotypes and WT in the reference (the top-left corner) are often absent from dbSNP [Sherry et al., 2001], while the tracking variants are almost always found in dbSNP, reinforcing our argument that the tracking variants are more likely to be TPs. Further, we found that the transition to transversion ratio (Ti/Tv) for the tracking variants was 2.15, which matches the expected ratio for humans (about 2.1). However, the Ti/Tv for the non-tracking variants was much lower (1.38); see Table S1.

To assess concordance we consider the 50Y/50C mixture since we should be able to detect all variants from both individuals. The union of the genotypes contained 5,230,349 alt allele calls. After restricting to regions with at least 20X coverage, 4,927,652 variants remained. Of those, 4,836,390 were called, for an FNR of 0.019. There were 5,605,654 variants called by VariantTools in the 50Y/50C mixture, of which 4,965,350 were also in the reference genotypes (TP). This yields an FDR of 0.11 assuming all predictions are wrong. However, we predicted genotypes for 5,346,889 of the variant calls, yielding a best case FDR of 0.028, assuming that our prediction algorithm correctly classified variants. This yields 381,539 new true positives. For the GATK calls, similar calculations estimate an FNR of 0.056 and an FDR of 0.005 (using the predicted genotypes from the GSNAP/VariantTools results). The GATK FDR is in agreement with the previously reported error rate of an error every 100-200kb [Li, 2014]. The higher FNR in GATK (0.056 vs. 0.019) suggests that GATK may be achieving a lower FDR by calling fewer positives. This strategy will result in more FNs.

### 4.4 Effects of homology and coverage

Figure 5B shows that the FDR is higher in homologous (self-chained) regions and also provides details on the relationship between FDR and coverage. Figures 2D and 2E reveal some of the differences in coverage that we obtain between the two pipelines and how self-chaining affects the coverage.

**Figure 5:**
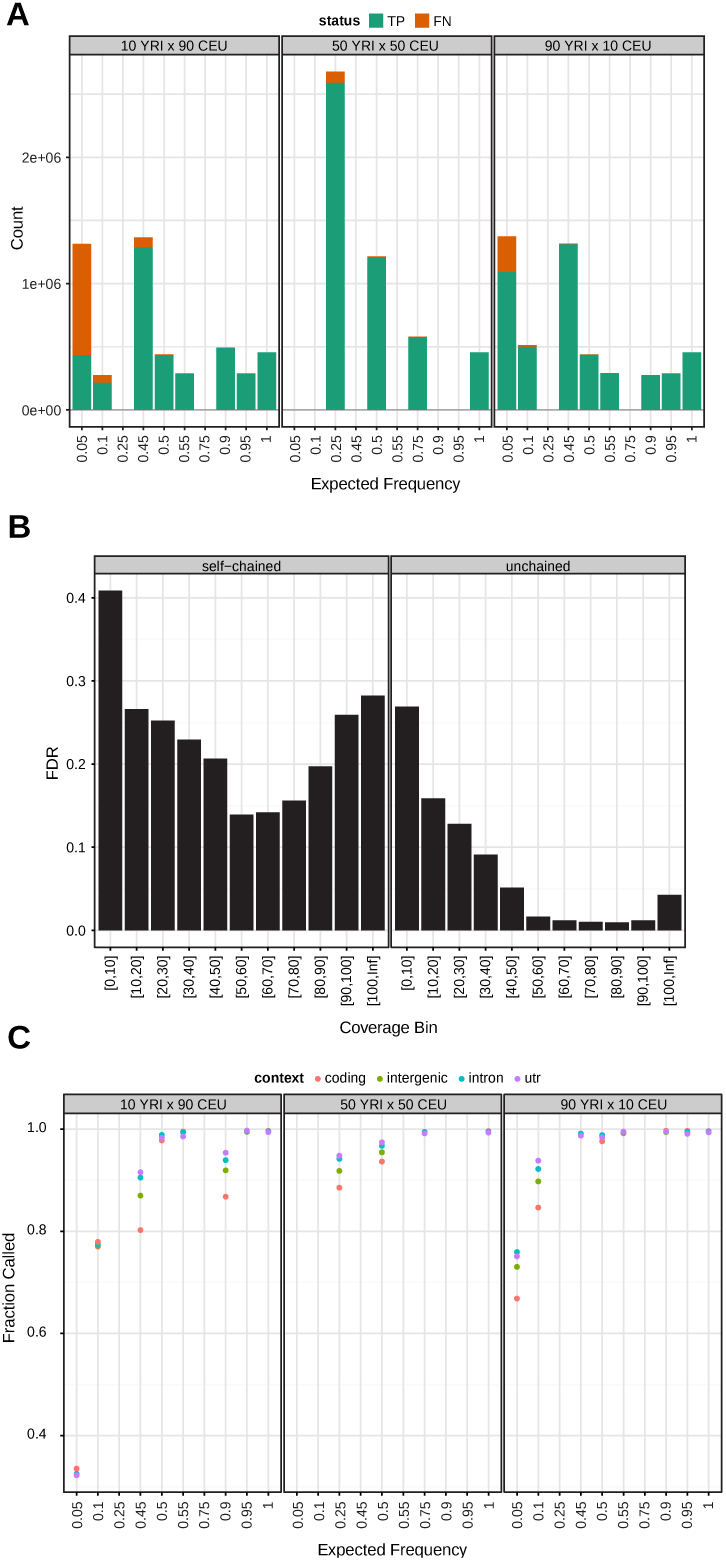
*Sensitivity and specificity.* (A) The count of variant calls from the reference genotypes at positions with >=20 coverage, conditioned on the expected frequency and whether the variant allele was called by the GSNAP/VariantTools pipeline (TP if called, FN if not). As expected, most of the FN are for variants expected at <=5% frequency. (B) FDR (counting tracking variants as TP, see Methods) for the GSNAP/VariantTools pipeline by coverage bin and self-chain status. The FDR is highest for low coverage regions and is also increased at extremely high coverage; the effect is magnified for self-chained regions, indicating that mapping is playing a role. (C) Fraction of the reference genotypes that GSNAP/VariantTools called; similar to panel A except conditioned on genomic context. The fraction is generally lowest in coding regions, perhaps because repeat masking rarely extends into coding regions.

High coverage may indicate that reads aligning to one locus in the reference originated from multiple loci in the individual, which can yield erroneous variant calls [Li, 2014]. Or the aligner may identify some reads as multi-mapping, and as these are often dealt with in an *ad hoc* manner the effect on frequency estimates is hard to predict.

Improper alignment may lead to clustered sets of variants. We found that high variant density is associated with self-chain status (see S9), and Figure 2C shows that false positive variants from the GSNAP/VariantTools pipeline have more neighbors than true positive variants. This motivated the variant proximity filter (see S1.3).

### 4.5 Effects of copy number variation

Copy number variation can affect the observed variant frequency and coverage. Since the probability of capture of the DNA is proportional to abundance, increases in copy number should result in increased coverage, while decreases in copy number should result in decreased coverage.

We obtained PCR-validated deletion calls from the HapMap project [Mills et al., 2011]. There were 190 CEU autosomal deletions that were validated by both the LSU and WTSI groups. They covered a total of 213kb and 92 unique YRI-specific variants. Figure 3C shows that the observed frequencies for YRI variants within CEU deletions are higher since the CEU deletions yield a higher proportion of YRI reads. We observe that the coverage at those positions decreases with decreasing YRI proportion, Figure 3D, as expected since there should be no contribution from the CEU. Figure S8 shows the coverage across mixtures for a single CEU deletion.

### 4.6 Sensitivity and specificity

Figure 5A shows the false negative rate (FNR) at each expected frequency for the GSNAP/VariantTools calls. As expected, the FNR is significantly higher for the 0.05 and 0.10 variants.

We also see an unexpectedly high FNR at 0.45 in the 10Y/90C mixture, and at 0.25 in the 50Y/50C mixture. Many of these positions lack coverage using our GSNAP pipeline as reads that align there were identified as multimapping. Indeed, about 58% of the FNs have a read-pair-concordant multimapping coverage at or above 10X, compared to 8% for the TPs.

We found that both the FDR and FNR depend on the genomic context, and, somewhat surprisingly, the FNR for the 50Y50C sample (Figure 5C) was highest within coding regions. Using Homologene paralogs [Coordinators, 2014] we found that the FDR (assuming that tracking variants are TPs) is higher in coding regions with paralogs (0.19, 136/437) than in those without (0.03, 1779/36191). Thus, accounting for homology can reduce the FDR in the coding regions to a value below the (unadjusted) FDR of the UTRs.

### 4.7 Validation of discordant calls

We believe that many of the putative FPs are actually real variants that were not detected previously and hence are not FPs, but rather TPs. We have shown that many of these variants have observed frequencies that track with the mixtures and are registered with dbSNP. Hence, they are consistent with being TPs. We believe that the tracking variants (where we predicted a genotype) will validate at a higher rate than those where we were unable to call a genotype.

For validation we selected variants that were in coding exons, not in selfchain regions and where the coverage was between 40X and 80X. Validation was carried out using the Sequenom technology. We selected 130 FP sites (86 tracking, 44 non-tracking). With Sequenom we were able to call a genotype for 106 of the sites (24 failed primer design). We observed a high validation rate, 66/68, for tracking variants, whereas only 10/38 of the non-tracking variants validate.

We also selected 137 TP sites where the predicted genotype did not match the published genotype. We were able to design primers for 129 sites, of which 84 were tracking and 45 were non-tracking. A total of 90 sites were published as unique to the CEU, and 37 as unique to YRI. We validated them with Sequenom. For the tracking variants, we found a concordance rate of 63/84 between our predictions and the Sequenom calls. Almost half of the errors (10/21) were predicted 0.5/0 but validated as 1/0. We may have underestimated the YRI frequency due to a bias against YRI in the actual mixture ratios (Figure 1). Of the non-tracking candidates, 10/24 were validated as wildtype, suggesting that some of the published calls are real, even when their frequencies do not track the expected trend. For more details on the validation, see S10.

## 5 Discussion

We have created a valuable data set that can be used to address the issues of accurate identification of point variants and their frequency. It also represents an opportunity to improve and enhance the development cycle for methodological advances. The existence of high quality *reference* data sets allows researchers to directly compare their approaches on identical data and provided (as is the case with our experiment) the data are sufficiently extensive these comparisons are valid. The effect of such a data set often results in both a convergence of algorithms on standard best practices and the clear elucidation of problems that have not yet been addressed.

We used these data to show that one can detect variants at low frequency without sacrificing specificity. We called 79% of the 10% variants in the 10Y/90C sample, while the overall FDR was only 0.028 in the 50Y/50C sample based on the validated assumption that the tracking variants are actually TPs. The primary calling filter considers the alternate read frequency, which we have shown is an accurate estimate of the underlying frequency. The performance of the algorithm benefits from exclusion of homopolymers and low complexity regions. However, while computational filters can overcome shortcomings in technology, there is a limit to their effectiveness. For example, we know self-chained regions are prone to error but excluding them will degrade sensitivity.

Further we showed that FDR was dependent on genomic context and showed that within coding regions homology was a major factor behind an increased FDR. We also showed that while others have reported extra-Binomial variation the phenomenon seems to be largely related to variants in regions with positive self-chain scores suggesting it arises due to mapping/alignment problems. Hence we suggest that the need for additional modeling complexity may be alleviated instead by accounting for the quality of the alignment.

While we agree with [Cibulskis et al., 2013] that the accurate detection of point substitutions is a key step in understanding the cancer genome, we believe that one has to go even further. In order to achieve the objectives of [Nik-Zainal et al., 2014] and provide researchers the ability to interpret the evolutionary history of the tumor we must not just identify the point mutants, but also characterize their frequency. Our analysis shows that we are able to accurately estimate the underlying allele frequency.

